# Aggregation dynamics of charged peptides in water: effect of salt concentration

**DOI:** 10.1101/649004

**Authors:** Susmita Ghosh, T Devanand, Upayan Baul, Satyavani Vemparala

**Affiliations:** The Institute of Mathematical Sciences, C.I.T. Campus, Taramani, Chennai 600113, India; Homi Bhabha National Institute, Training School Complex, Anushakti Nagar, Mumbai, 400094, India; Institue of Physics, Albert-Ludwigs-University of Freiburg, Hermann-Herder-Strasse 3, 79104 Freiburg, Germany

## Abstract

Extensive molecular dynamics simulations have been employed to probe the effects of salts on the kinetics and dynamics of early-stage aggregated structures of steric zipper peptides in water. The simulations reveal that the chemical identity and valency of cation in the salt play a crucial roles in aggregate morphology of the peptides. Sodium ions induce the most aggregated structures but this is not replicated by potassium ions which are also monovalent. Divalent Magnesium ions induce aggregation, but to a lesser extent than that of sodium and their interactions with the charged peptides are also significantly different. The aggregate morphology in the presence of monovalent sodium ions is a compact structure with interpenetrating peptides, which differs from the more loosely connected peptides in the presence of either potassium or magnesium ions. The different ways in which the cations effectively renormalize the charges of peptides is suggested to be the cause of the differential effects of different salts studied here. These simulations underscore the importance of understanding both the valency and nature of of salts in biologically relevant aggregated structures.

## I. INTRODUCTION

The process of aggregation of proteins in solution has been implicated in a wide range of diseases including Type II diabetes, Alzheimer’s, Parkinson’s and Huntington’s disease^1,2^. Understanding the underlying causes for the formation of aggregates of proteins, which are otherwise individually solvable, is far from complete. Given the important role of such protein aggregates in the aforementioned neurodegenerative diseases, it is also crucial to examine the role played by solvent conditions, including pH, temperature, salt content etc., on promotion or degradation of such protein aggregates^3–5^. Much of the focus in protein aggregation studies has been on the formation of fibrillar structures, like in the case of amyloid aggregates, since they are directly related to diseases^6,7^. Proteins, on the other hand, are also known to form amorphous aggregates which may lack precise structural information, but are nevertheless not the desired end product for protein solubility and protein functional aspects^8–11^.

The phase separation behaviour of proteins is strongly linked to the protein sequence along with the solution conditions, and is driven by complicated balance of interprotein, intra-protein, protein-solvent, solvent-solvent interactions along with the entropic contributions. Notably, proteins containing the so called “low-complexity regions”(LCRs) have the propensity to form condensates and the propensity is modulated by conditions such as temperature and ionic strength as reviewed in the past^12,13^. Transitions from individually collapsed states to aggregate phase of LCRs are mainly demarcated by hydrophobicity, aromatic and aliphatic character, and charge content in low-complexity sequences^14–22^. The phase transition of proteins enriched with hydrophobic amino acids is likely to be mediated by hydrophobic interaction under physiological conditions and they are aggregation prone in general^23^. In contrast, LCRs with charged residues would result in high solubility and might prevent aggregation due to unfavorable electrostatic interaction between charged residues^24,25^. Understanding the molecular basis of such underlying phase transitions of charged LCRs at varying physiologically conditions is of particular interest.

Aggregation behavior of charged proteins/peptides considered as charged biopolymers, having a coarse-graining description have been studied rigorously before^26–41^. In such simulations, for simplicity, polymers are assumed to be fully charged and aspects of their single chain behavior (aspects of compaction) and multiple chain behavior (aspects of aggregation) in the presence of counterions have been explored. As salt occurs universally in physiological environment, there is an increasing need to understand the role played by solvent, specific nature of salt or counterions in the solution and the structural conformational entropy of biological charged polymers themselves^42–47^.

The presence of salt in the process of protein aggregation has attracted wide attention because of its profound effect on protein aggregation, protein stability and protein solubility^48–54^. Previous studies have explained the effect of salt on protein aggregation in terms of various models such as the Debye-Huckel screening of charges, effects on the protein-water-ion interactions (Hofmeister series) and specific ion binding (electroselectivity series). Some of the parameters such as pH, temperature, presence of co-solutes etc., have been shown to influence the self-assembly of proteins in diverse ways as reported in several studies. Recently the experimental kinetic studies of salt-induced aggregation on a model antibody using different types of salts at various pH values have shown that salt concentration promotes aggregation through the formation of protein intermediates characterized by partially ordered secondary structures and is strongly dependent on ion’s specificity and pH conditions^54^. In the experimental studies on the yeast prion protein Sup35, salts were found to follow Hofmeister series in modulating the aggregation propensity^53^whereas specific anion binding was shown to play a crucial role in the aggregation process of mouse prion protein and b2-microglobulin^48,55^. In the case of Alzheimer’s Aβ(1-40) peptide both of the Hofmeister effect and anion binding effect play an important role on aggregation propensity and the aggregation was found to be progressively more favourable as the salt concentration was increased from 50mM to 500mM^52^. The heat induced aggregation kinetics of Human carbonic anhydrase II was also found to be highly sensitive to salt concentration^49^. There is also a report where different Hofmeister sequences are effective in the different concentration regimes of the salts^56^.

In this work, our primary objective is to probe the effect of different salts at moderately high concentration, and their valency on the aggregation of an explicitly solvated charged polypeptide system. In the present study, the aggregation dynamics of a hexapeptide (^57,58^ VEALYL, referred henceforth as IB12), a segment from the B chain of the fibril-forming protein insulin, was explored using atomistic molecular dynamics. This particular short peptide was selected in our studies of probing atomistically aggregation process for computational feasibility and also, it has been investigated earlier as a model system for protein aggregation studies^59,60^. Insulin, a small hormone consisting of α-helix and cross-β structure has become a model peptide for studying fibril formation because of it’s simple structure. Under certain conditions (elevated temperatures, low pH, hydrophobic interface, ionic strength, and mechanical agitation)^61,62^ it undergoes denaturation and further aggregates in fibril-like structures via fibrillation. Previous simulations^59,60^ on IB12 peptide using GROMOS force-field parameters reported an abundance of *β*-sheet rich aggregate formation from unstructured monomers and so, those studies are primarily focused on the initial aggregation events of fiber formation (natural or amyloid). However, from the oligomerization of IB12 peptide sequences, it is becoming increasingly apparent that it may be the prefibrillar (often amorphous and granular) aggregates that produce cytotoxic effects in vivo^8–11^. These require detailed atomistic simulations with correctly parameterised forcefields to capture the essential aspects of the proteins, water and salts added and can be very expensive computationally to explore the full aggregation pathways. It is more feasible, at atomistic resolution, to explore the initial pathways of aggregation, from which valuable insights into the early stage kinetics of aggregation can also be gained^59,60,63,64^.

Here we focus on the detailed understanding of the pathways of formation (and degradation) of amorphous aggregates, which are likely to be fundamentally different from the mechanisms of fibril growth. Several chains of IB12 (net charge=-1) peptide system were dispersed into three types salts namely NaCl, KCl, and MgCl_2_ solution at 1.0M concentration and the role of salt condition in accelerating and/or retarding the amorphous aggregation is discussed here. Attempts have been made to identify specific molecular interactions that direct the association of IB12 peptide monomers leading to the formation of stable aggregates in the various salt-solutions at molar level concentration. Our results provide mechanistic insight into the underlying physical processes of amorphous aggregation and explain the dependence of aggregation properties on the nature of salt by the detailed analyses involving Molecular Mechanics Generlised Born Surface Area (MM/GBSA) model, Charge renormalizatin of aggregates by salt-ions and Markov State Model (MSM). The MM/GBSA model has been used successfully in several areas of protein biophysics such as binding affinities protein-ligand, protein-protein and multi-component protein interaction^65,66^. MSMs^67–71^ are generally used in kinetic studies of folding and unfolding events of nucleic acids and proteins^72^. The charge renormalization of highly charged macroions has been observed when the macrions undergo a condensation in solutions in the presence of multivalent counterions^73–75^. Here the combination of MM/GBSA, MSM and Charge renormalization method will be important to protein aggregation studies as well.

The rest of the article arranged as follows: in section II we discuss the procedure to set up the systems, simulation protocols employed and methods used for our calculation. The results obtained are presented in section III, followed by discussions on the important findings of this study and the conclusions in section IV.

## II. METHODS

Aggregation of peptides IB12(Val-Glu-Ala-Leu-Tyr-Leu)^59,60^ is studied using all-atom classical molecular dynamics methods. Several IB12-solvated systems were prepared with 1.0M concentration of salts (both monovalent and divalent) and TIP3P^76^ water model. TIP4P-Ew^77^ water model was also used to analyse if the aggregation behavior depends strongly on the water model used.

All results in the paper are for TIP3P water model and the results from TIP4P-Ew are available in Sec. I of supplementary material. For ions, standard CHARMM parameters used extensively in biomolecular systems with TIP3P water model^78^ and similar parameters specific to TIP4P-Ew water model were used^79,80^. Each IB12 peptide carries a total charge of −1 due to the presence of Glu amino acid.

### A. System setup

The starting configuration of IB12 peptide structure was generated using *molefacture* plugin in VMD^81^ and the single IB12 peptide was solvated in TIP3P water box(≈ 38.5Å×38.5Å×38.5Å) and neutralized by adding a single Na^+^ ion in the water box. Initially the system was relaxed using conjugate-gradient energy minimization algorithm by 1000 steps and equilibrated for 5ns under isothermal-isobaric (NPT) ensemble condition at pressure of 1 atm with a timestep of 2fs. Next, 64 replicas of equally spaced equilibrated IB12 peptides were solvated in a cubic box of size 108Å^3^. Multiple systems were derived from this primary one by ionizing with 1.0M concentration of NaCl, KCl and MgCl_2_ salts using *autoionize* plugin of VMD. A separate control system was also prepared with addition of ions only to neutralize the total peptide charge. The systems were then equilibrated with 1fs timestep for 5ns under NPT ensemble by fixing the center of mass locations of the Ala residue in each peptides so that initial aggregation is prevented, allowing water and ions to equilibrate around the peptides. The final conformation for each case, at the end of this 5ns of equilibration run was used as initial system configuration for the subsequent production (NVT) runs. Two independent 100ns long simulations were performed for each case. All the system and simulation details are listed in Table I.

**TABLE I.** System details and summary of performed simulations for IB12 peptides in TIP3P water model.

| salt | salt conc. | No of cations | No of anions ($\text{Cl}^-$) | time(ns/run) | no. of sim |
| --- | --- | --- | --- | --- | --- |
| NaCl | 1.0M | 675 | 739 | 100 ns | 2 |
| $\text{MgCl}_2$ | 1.0M | 707 | 1350 | 100 ns | 2 |
| KCl | 1.0M | 675 | 675 | 100 ns | 2 |

### B. Simulation protocols

All MD simulations were performed using simulation package NAMD 2.9^82^ using CHARMM36^83^ parameters for peptides. The MD simulations were performed at a temperature of 298K in NVT ensemble. For NPT runs, a pressure of 1 atm was maintained using Nosé-Hoover-Langevin piston^84^ with a decay period of 100fs and a damping time of 50fs. The non-bonded cut-off distance was set to 12Å with a switching distance between 10Å and 12Å. The long-range electrostatic interactions were treated by Particle mesh Ewald (PME)^85^ method and the data visualization and analyses were done using the software VMD^81^ and in house data analysis scripts.

### C. Markov state model analysis

To investigate the time-evolution of aggregate size distribution Markov State Model (MSM) approach was used for analyzing the MD trajectories. MSM provides a convenient way to model kinetic network for conformational transitions^67–71^. In MSM formalism, the dynamics of the system is described in terms of state-to-state transitions and the time-evolution of the system is governed by the discrete-time master equation:

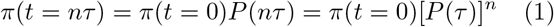

where *π* is the row vector of probabilities of occupying any of the Markov states at time *t* and *P*(*τ*) is the transition probability matrix, whose entries, *P_ij_*, provide the probability of the system to be found in state *j* at time *t* + *τ* given that it was in state *i* at time *t*. *τ* is the time interval of obersevation, called the lag time.

Here, MSM describes the dynamics among the aggregate states using a transition matrix, *T*. The components of *T*, *T_ij_* were computed by counting the total number peptides transitioning from the aggregate state of *i*-mer (*S_i,t_*) to aggregate state of *j*-mer (*S*_*j,t*+1_) in next time step. Here *S_i,t_* and *S*_*j,t*+1_ represent an aggregate size state of *i*-mer and *j*-mer at time *t* and *t* + 1 respectively. A 64 × 64 transition probability matrix, *P* was constructed by normalizing the transition matrix elements with the sum of elements contained in the corresponding row. Mathametically we can express this term as follows;

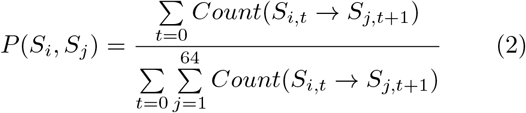

Thus, the fraction of aggregate size (*N*-mer) in the trajectory after *n* propgation steps can be obtained by solving the master equation as row vector *π*(*n*) = *π*(0)*P_n_*. Here *π*(0) is the row vector containing the starting fractional populations as *π*{1,2,.…, 64;t=0}={1,0,.…,0}.

## III. RESULTS

### A. Aggregate morphology

In this section the dependence of the size of the peptide aggregates and their morphology on the concentration and valency of the salts are described. At any given instant, two peptides are defined to be part of the same aggregate if the distance between two non-H atoms in the respective peptides is less than 3Å. The representative snapshots of aggregate states for all three salts under 1.0M concentration and the control simulaton at the end of 100ns, are shown in Figure 1. The individual peptides are colored based on the size of the aggregate they belong to. From the Figure 1a, it can be seen that for systems with NaCl salt, the number of aggregates is smallest, with one single aggregate being the predominant aggregate morphology. This is in contrast to the control system (Figure 1d), where the number of aggregates is larger and the average aggregate size is much smaller. The typical aggregate morphologies in other salt systems is also shown in the Figure 1. In the case of KCl (Figure 1c), also a monovalent salt, the aggregation behavior is very different from that of NaCl case, strongly underscoring the influence of type of cation on such aggregation processes and not just the valency of the cation. In the case of divalent salt MgCl_2_ (Figure 1b), it can be seen that the though aggregate size is larger than KCl and control systems, the system does not display the near phase separated aggregate morphology seen in NaCl systems. These visual results suggest that the aggregate behavior of peptides in salt-water systems are function of not only the valency of the cation, but are strongly dependent on the cation type as well.

**FIG. 1.**
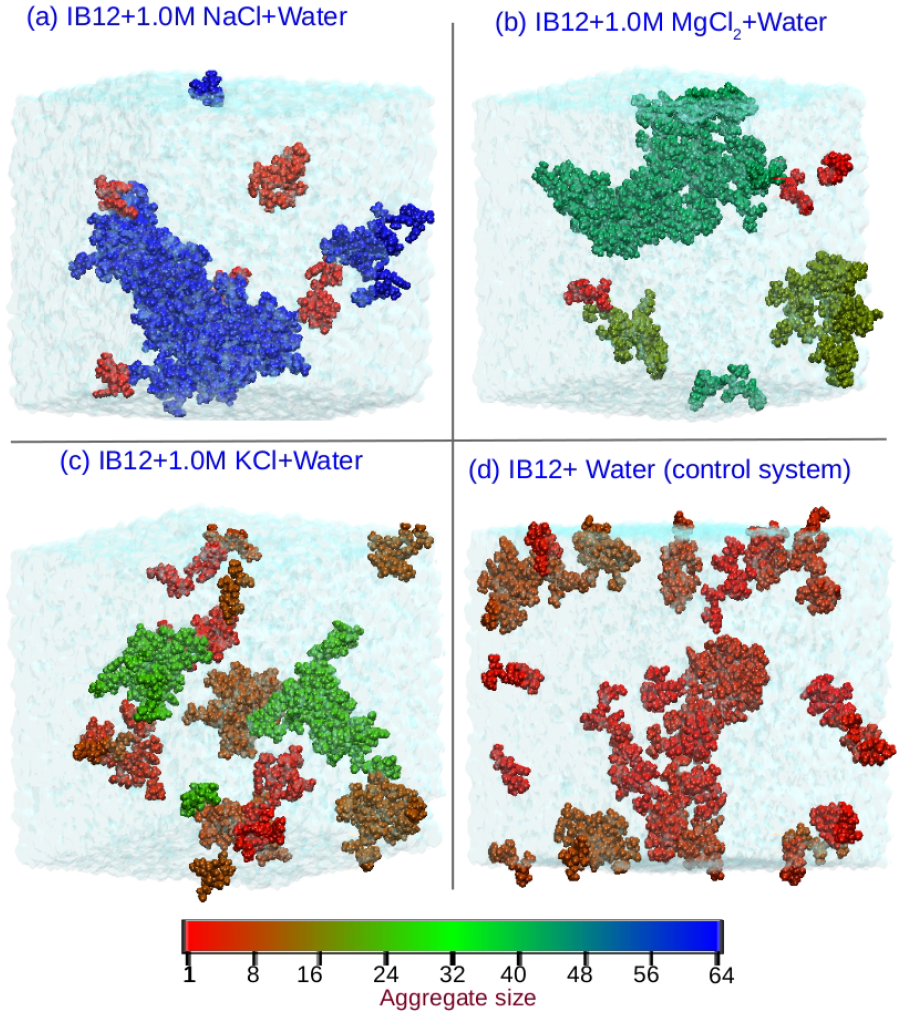
MD snapshots (in VDW representation) of the end-product in one of the 100ns long IB12 peptide simulations for NaCl(a), MgCl_2_(b), KCl(c) salt and control system(d) respectively. The coloring of the peptide is based on the aggregate size they belong to. The salt-ions are not shown for clarity.

To quantify the visual results in Figure 1, the morphology of the aggregates is quantified in terms of normalized aggregate size distribution, over the last 10ns, which for all the systems is shown in Figure 2. In Figure 2 the aggregate size (*S*) is the total number of peptides forming a given aggregate. Apart from a few stray single peptides, the distribution of the aggregate size for 1.0M NaCl salt solution, peaks around ~58 at the end of 100ns long simulation suggesting that for this case, almost all of the peptides are in an isolated aggregate. The distribution of aggregate sizes for KCl and MgCl2 suggests that the propensity in these systems is not towards one single large aggregate, but a distribution of aggregate sizes corresponding to the well known finite bundle model suggested in literature^86–89^. In finite bundle model, the finite-sized bundle-like aggregates of the charged macromolecular system in the presence of multivalent salt are suggested to be the thermodynamic equilibrium states. However, in the control system where no extra salt is added, the distribution of aggregate sizes is restricted to smaller sizes, clearly showing that the effect of addition of salt, in general, is to promote the aggregation though differences may exist in the morphologies of the same depending on the valency and type of the cation. The increase of aggregate sizes in the presence of NaCl salt, compared to control system seen in our simulations agrees with similar increase in aggregate sizes seen in experiments^90^ on chitosan based charged polymers. These experiments also suggested that the aggregation morphologies in presence of divalent salts is different from that of monovalent salts.

**FIG. 2.**
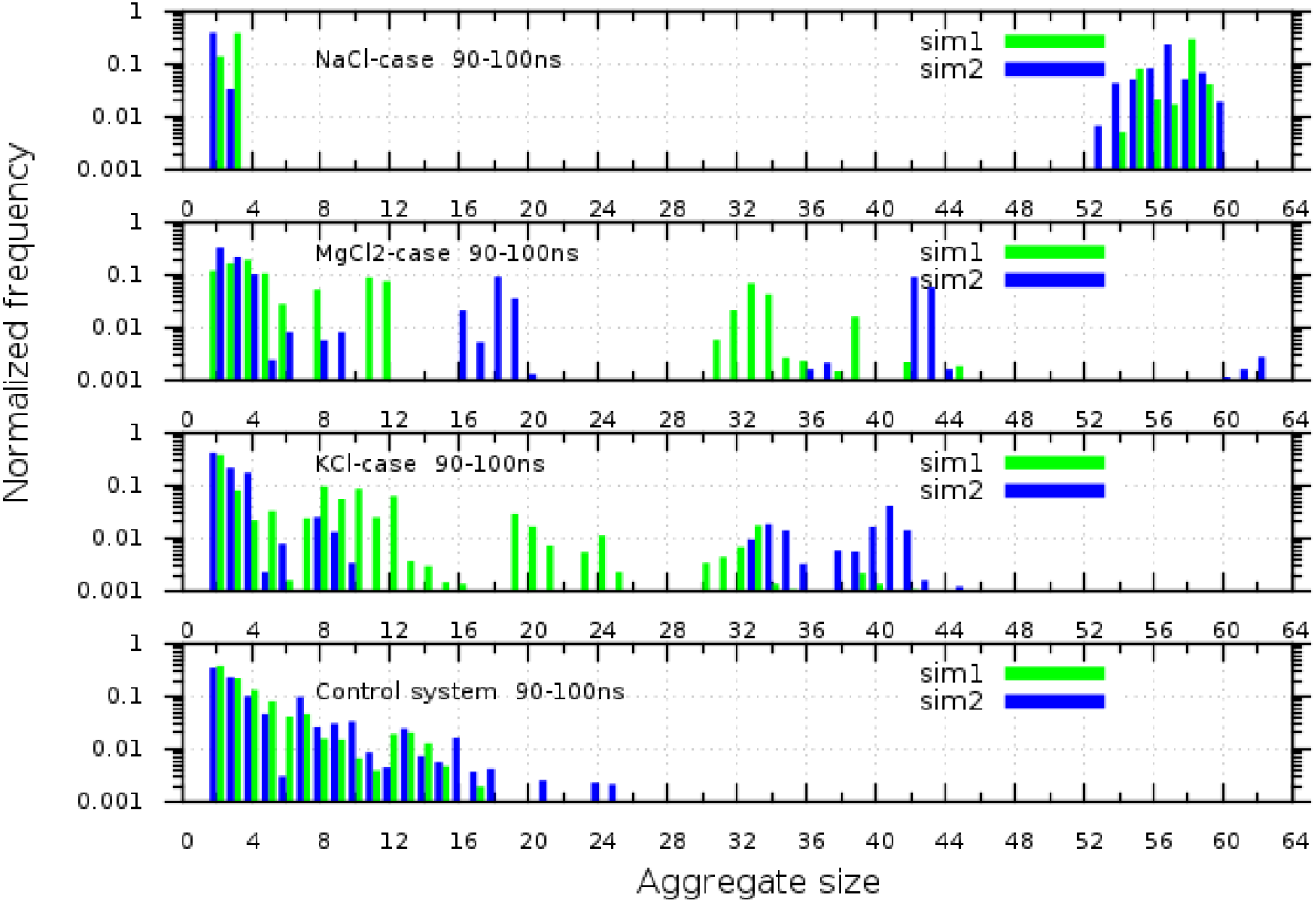
Normalized aggregate size distribution for IB12 system with NaCl, MgCl_2_, KCl salt-solutions and control system computed from last 10ns simulation data for each trajectories.

One of the measures of aggregate size is radius of gyration (*R*_g_) and particularly its variation with time. In Figure 3, *R*_g_ of the largest aggregate is plotted as a function of time in the last 25ns of simulation to understand the differences in the size distribution of the aggregates between NaCl and KCl systems though in both systems the cation is monovalent. The data in Figure 3 shows that even though the aggregate size for KCl solution is much smaller than that of NaCl, the *R*_g_ of the largest cluster for KCl (33 peptides) is similar or larger than that of NaCl case (58 peptides) indicating the possibility of more compact structure formation of the peptides in the presence of NaCl salt. This can also be visually seen from Figure 1c, where the aggregates, in the presence of KCl, are more loosely packed.

**FIG. 3.**
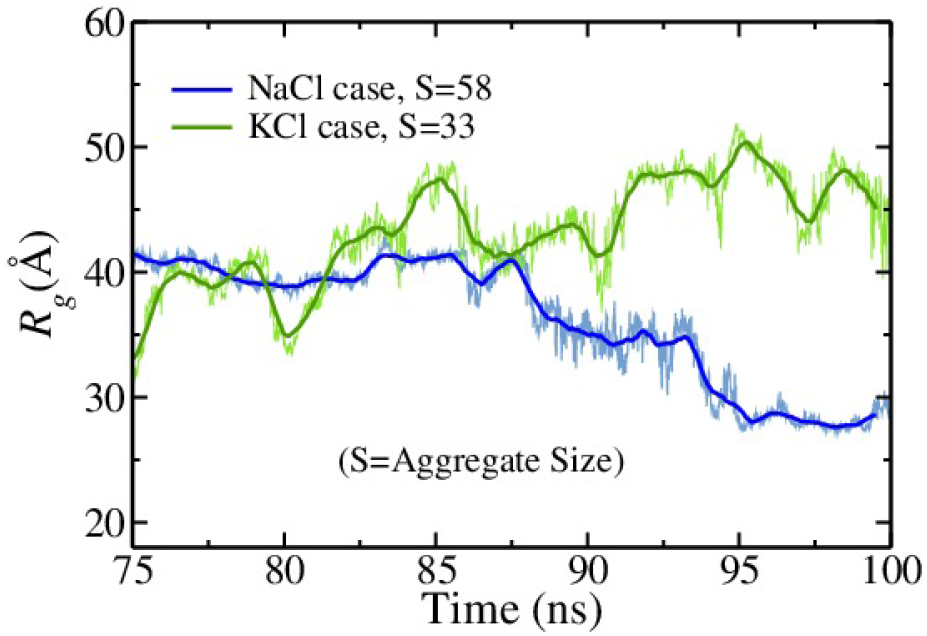
Time evolution of radius of gyration (*R*_g_) of large peptide aggregate for IB12-NaCl and IB12-KCl systems in the last 25ns of simulation. *R*_g_ values are calculated for the peptide chains belong to aggregate of size 58 and 33 for NaCl salt and KCl salt solutions respectively.

To quantify the compactness of the aggregates, we compute the pair distribution function g(r) for hydrophobic residues of the peptides as shown in Figure 4. The peptide composition, as seen in Methods, shows that there is a single charged amino acid GLU and has three hydrophobic residues (VAL, ALA, LEU). From the g(r) data, it can be seen that the location and peak heights for ALA-ALA distribution is at a shorter distance for NaCl than for KCl. This can be attributed due to different sizes of the cations. NaCl promotes the aggregation of hydrophobic residues much more compared to the KCl and the most probable closest distance between hydrophobic residues is much larger for the case of divalent salt MgCl_2_. Some of the earlier simulations on small hydrophobic solutes and hydrophobic polymers in salt solutions have shown strong stabilization of these hydrophobic interactions in the presence of NaCl^91^. These observations strongly suggest that valency and nature of salt play a crucial role on the morphology of the aggregates and the aggregate sizes. A possible understanding of dependence of critical size of aggregates and the valency of the ions is discussed in latter sections.

**FIG. 4.**
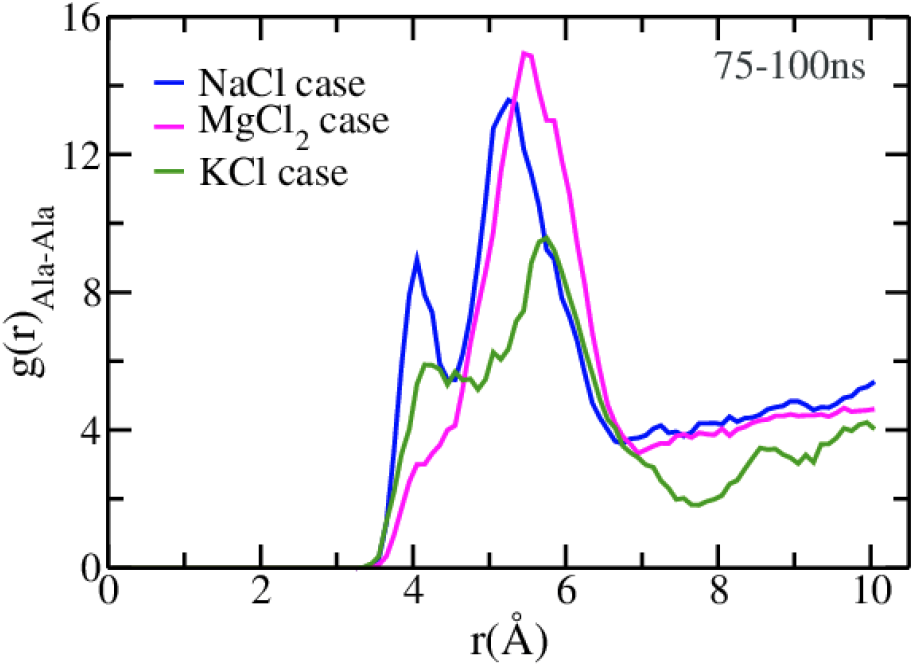
The pair correlation function (g(r)) of the hydrophobic residues Ala of the IB12 peptide for three types of salt solutions.

A visual inspection of the IB12 simulation trajectories suggested that there was a conversion from random coil to extended *β*-sheet conformation as the simulation time progresses which is generally in line with earlier literature^59^. In a couple of cases, there were signatures of *β*-sheet formation between two peptides but they were not so substantial. Representative snapshots of secondary structures from one of the IB12 simulations are shown in Figure 5 for NaCl and MgCl_2_ salt solutions. For both cases very few monomers (~4-8 strands) were found to form parallel, anti-parallel or disordered beta-sheet conformation. In any case, *β*-sheet formations are expected to take place at considerably longer timescales. However, in the present study we are interested in the early stage of salt-induced aggregation events of charged peptides rather than conformational dynamics.

**FIG. 5.**
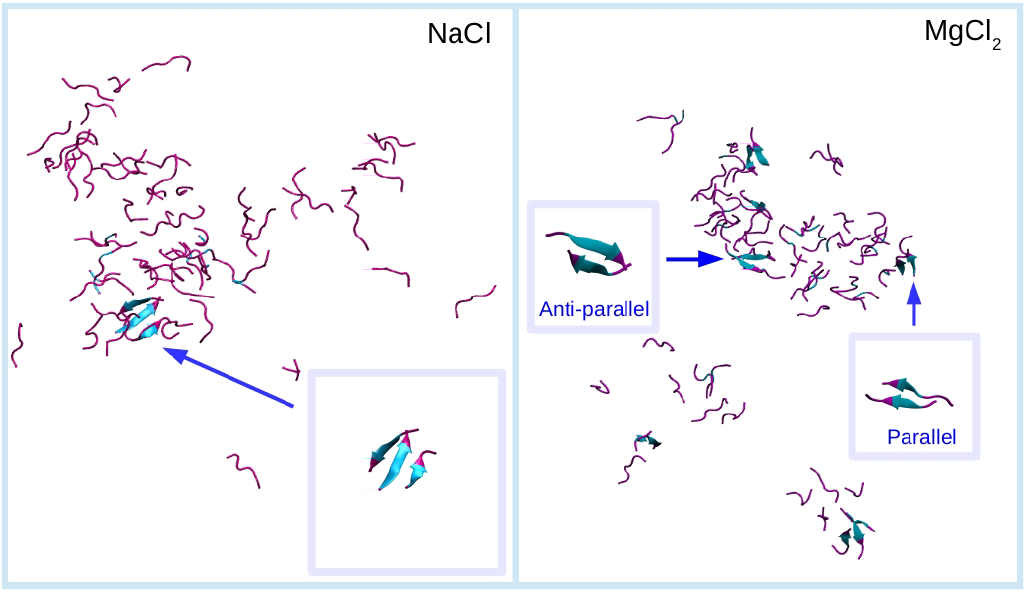
Snapshots of secondary structures observed at the end of 100ns long trajectory of IB12 in NaCl and MgCl_2_ salt solution. Two set of representative peptide aggregates end-structure (at 100ns) are depicted in cartoon representation, showing ordered or disordered *β*-sheet assemblies. The colors encode *β*-sheet (cyan) and random coil (magenta) secondary structure elements.

To understand the effect of salt on the dynamics of peptide aggregation at molecular level, several comparative analyses between different kind of IB12-salts and control system have been done. In next section we describe the dynamic evolution of peptide aggregates.

### B. Dynamic evolution of peptide aggregates

Figure 6(a) presents a comparison between the evolution of the total the number of aggregates in the four systems. In Figure 6(a), *N*(*t*) is the the number of aggregates at time *t* with aggregate size, *S* >=2. In complete phase separated state, *N*(*t*) = 1 *i.e*. one single aggregate containing 64 peptide chains and in complete deaggregated state, it is 0. For finite bundle^86–89^ *N*(*t*) lies somewhere in between 1-32 for 64-mer peptide system. In plot Figure 6(a) we see that the peptides started to aggregate from very early stage of simulation for all four cases. The plot suggests that assembly towards the final oligomeric state is faster in the presence of salt as compared to control system. It is northworthy that aggregation is relativey faster for NaCl case than KCl salt at long-timescale. A slight difference in aggregation rate was observed between NaCl and MgCl_2_ salt beyond 60ns. The deaggregation events of IB12 multi-peptide system in MgCl_2_ at timescale between 60-80ns is accounting for this difference in aggregation rate. The time evolution of the fraction of peptide chain taking part in aggregate formation are shown in Figure 6(a) inset. Our result shows that almost 90% of peptide chains (out of 64) formed aggregate in the presence of salt whereas the fraction is around 80% for control system, which also has a propensity of forming substantially smaller aggregates.

**FIG. 6.**
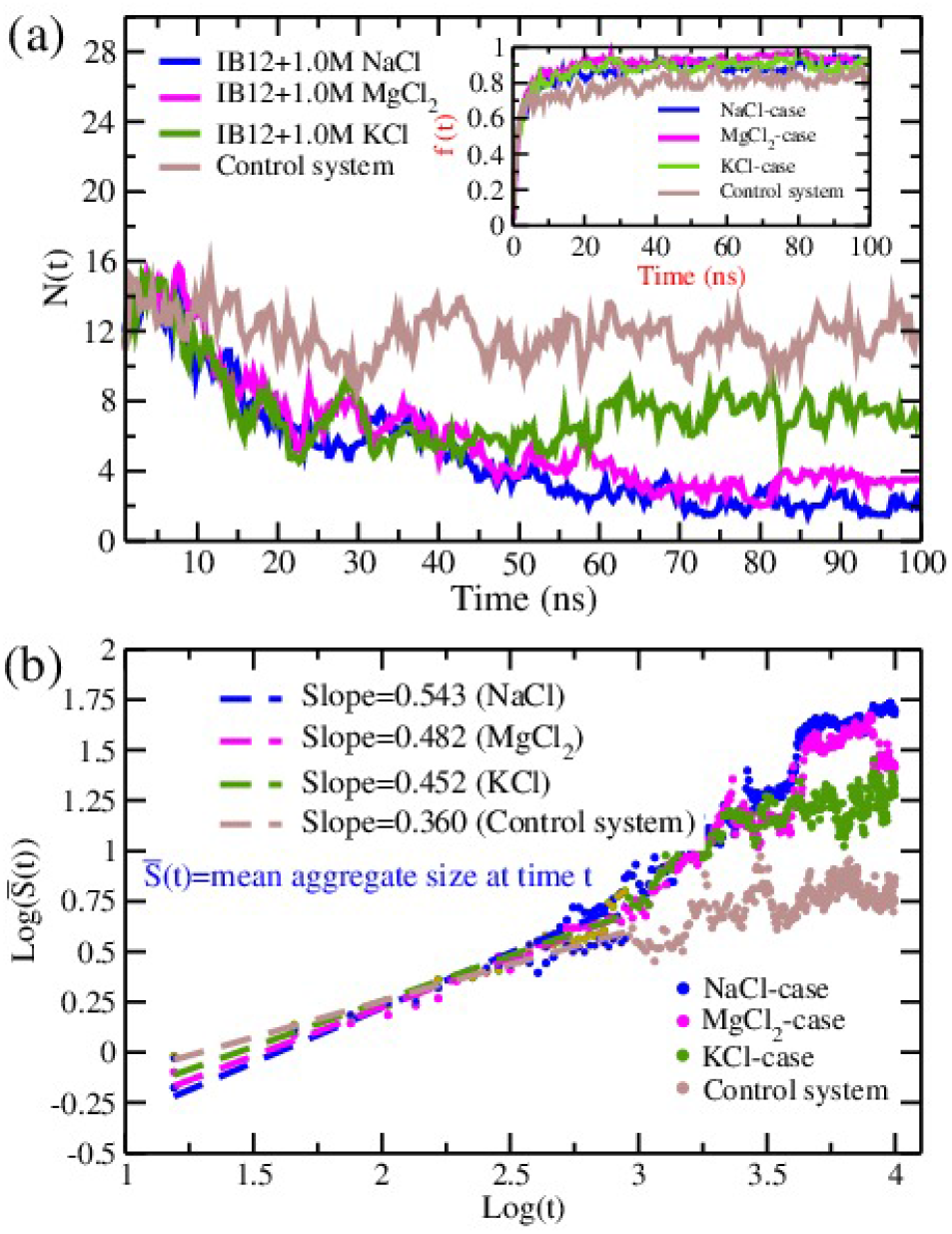
(a) Dynamic evolution of the total number of aggregates *N*(*t*) excluding single peptide state i.e. aggregate size (*S*)>= 2. The blockavearge of data were obtained from each time slice of 300ps. Inset shows the time evolution of the fraction of total no. of peptide chains forming aggregates. (b) Dynamic evolution of the mean aggregate size 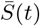 in the 64-peptide system. The dashed line, representing fit of 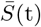 at timescale of 0-10ns. The slope of the straight lines are about 0.543, 0.482 and 0.452 for NaCl, MgCl_2_ and it is ~0.360 for control system respectively.

In Figure 6(a) we observe that initially the growth of aggregates size proceeds rapidly for all cases and beyond 75ns timescale the aggregate phase were emerged to be more stable. Large stable aggregates were formed at timescale of 75ns. The average number of aggregates formed at time scale of 75-100ns are 2.11, 3.34, 7.54, 11.738 for NaCl, MgCl_2_, KCl and control system repectively.

For 1.0M MgCl_2_ salt solution, at the timescale of 75ns a large cluster with aggregate size >60 was found to be formed in one of our 100ns trajectories but as the time progresses deaggregation took place and the larger cluster collapsed into two smaller clusters of sizes ~33 and ~18. The aggregate formation of the peptide system was found to be more stable in NaCl salt solution compared to MgCl_2_. We can define the mean aggregate size 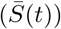 at time t by the standard formula used in aggregation of clusters (Refs.^92,93^)

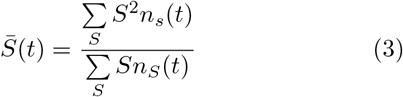

where *n_S_* is the number of aggregates (excluding single peptides) that contain S peptide chains. Figure 6(b) shows 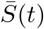 as a function of time. At short timescale, 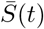 is found to grow with time as a power law but, the dynamic scaling for aggregation of clusters is not followed at long timescales. The lines, representing linear fit of 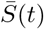 at short time, have a slope of about 0.543, 0.482 and 0.452 for NaCl, MgCl_2_ and it is ~0.360 for control system.

To monitor the deaggregation events in salt solution we put our focus on the aggregate state (*S_i,t_*) associated with each peptide chain at each time step t which are represented by Figure 7. Figure 7 depicts the time series of aggergate size states associated with each peptide for NaCl and MgCl_2_ salt along y axis, where the x axis labels the peptide segment running from S1 to S64 and the colorscale being a mark of aggregate size state (*S_i_*) to which a particular peptide belongs. Here *S_i_* represents an aggregate size state of *i*-mer. In matrix representation of following figures, the decrease in intensity from blue to green in any column implies the occurance of disassociation event. At the early stage of aggregation the peptide molucules were found to be associating and disassociating reversibly, eventually large aggregates were formed between 75-80ns. Beyond 75ns timescales, deaggregation events have been observed predominantly for MgCl_2_ salt compared to two other salts. This result suggests us that the presence of divalent atom may have destabilising influence on the association events of peptide molecules. So an understading of the interaction between the protein, ion and water molecules can allow us to get better insight into the mechanism of aggregation process in the presence of salts, those are follwed in the succeeding sections.

**FIG. 7.**
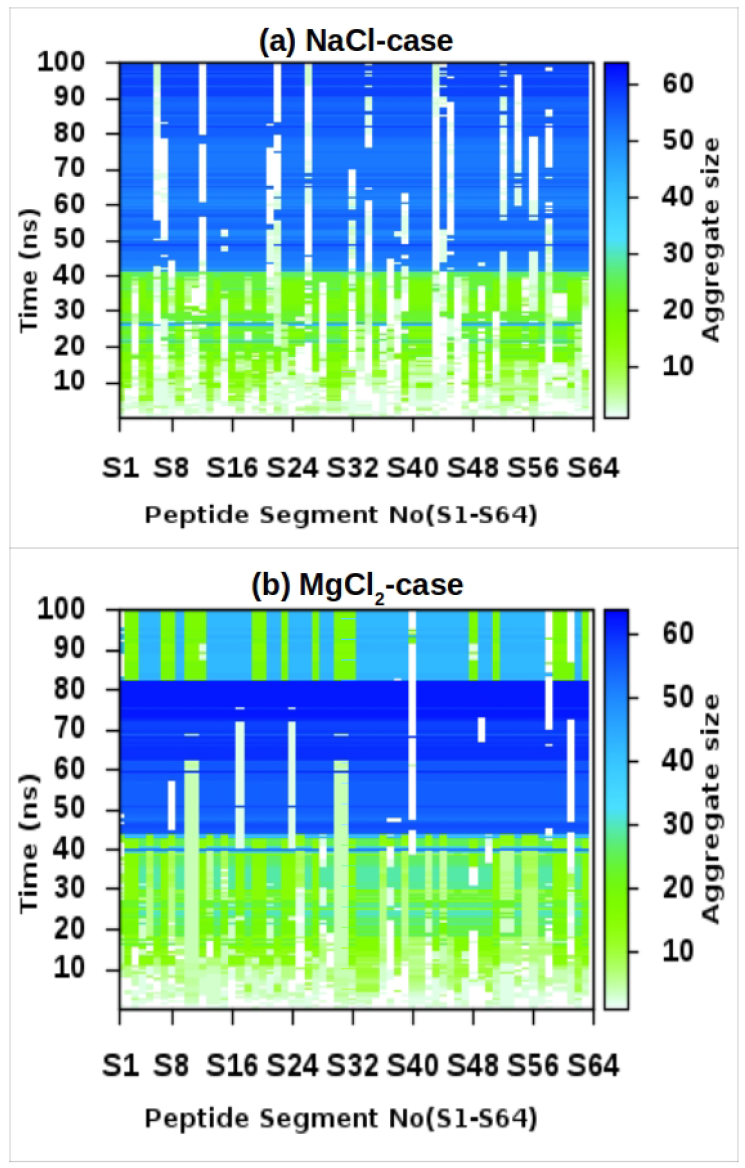
Matrix representation of the time evolution of aggregate size state (*S_i_*) of each peptide monomer for NaCl(a) and MgCl_2_(b) salt-solution. The x-axis denotes the labeling of peptide segments running from S1-S64 and the color scale encodes the aggregate size state *S_i_*.

### C. Interaction between water, salt-ions and peptides

To assess the interaction among the peptide, salt and water molecules we computed the radial distribution function, g(r) for various cases. The radial pair distribution function (pair correlation function) provides us valuable insights into the distribution of atoms (molecules) around another atom (molecules) of same kind or different kind. First we calculate the distribution of salt-cations around Glutamic acid (Glu) of peptide molecule from 75-100ns simulation data as shown in Figure 8.

**FIG. 8.**
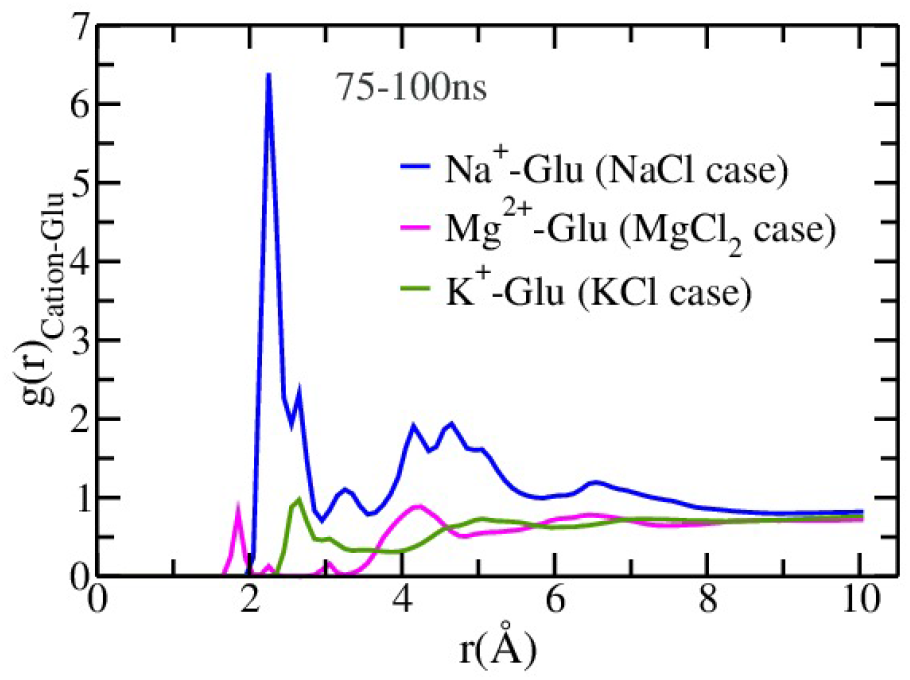
Radial pair distribution function (g(r)_cation-Giu_) of cations of salt around the negatively charged residue Glu of the peptide in salt-water solution obtained from last 25ns MD data.

The g(r) results clearly show that Na^+^ ions interact most strongly with Glu residues compared to both K^+^ and Mg^2+^ cations. This also indicates that the total number of cations within 3A of residue Glu in NaCl case is much higher compared to other two salts and also that more Na^+^ cations are interacting with Glu in comparison with K^+^ and Mg^2+^ ions. For MgCl_2_ salt though a peak appears at smaller distance compared to NaCl, the effective distance between a typical Mg^2+^ cation and Glu is much larger and is greater that 4A compared to the 2Å most probable distance between Na^+^ and Glu residues. The order of the strength of ion-pair interaction in aqueous solution of three salts was found as NaCl<KCl<MgCl_2_ from the calculation of g(r) of anions around cations (see Sec. II of the supplementary material). The interaction of salt-ions with water also plays a crucial role in solubility of a salt in water as the solubility is the result of the balance between two competing interactions: the ion-pair interaction of the salt and the ion-water interaction. From the g(r) analysis of water-cation interaction we observed a well-defined strong hydration shell around Mg^2+^ ions with an additional outer hydration layer whereas for monovalent salts, Na^+^ and K^+^ ions exhibit weaker interaction with water (see Sec. II of supplementary material). However, any effects of high salt concentration of salts on water’s bulk structure were not studied in our simulations. We noticed that within the hydration shell around Mg^2+^ the oxyzen atoms of hydrating water molecules pointing towards the Mg^+^ ion make relatively stronger interaction due to divalency and negative hydration entropies of smaller Mg^2+^ ions compared to realatively larger monovalent ions (the snapshots of condensed anions and water molecules around a single cation for three types of salt-solutions are shown in Sec. II of supplementary material). These results are consistent with the Hofmeister series ranking chaotropes (weakly hydrated) to kosmotropes (strongly hydrated) for cations as NH4^+^>Cs^+^>Rb^+^>K^+^>Na^+^>Li^+^>Ca^2+^>Mg^2+^^94–97^. It is conceivable that in the case of MgCl_2_ salt, the water molecules within the first hydration shell of Mg^2+^ ion are so tightly bound that Mg^2+^ ions are less available for peptide atoms.

Also, other factors like the charge, size, polarizability, temperature, pressure and concentration of the ions contribute to the the solvent dynamics, mobility and solulibility of salts. The vdW radii of three metal ions studied here are 2.27Å, 1.73^Å^, 2.75A for Na, Mg and K atom respectively^98^. Now, we define the ions/atoms of type A which interacts with another atom/molecule of type B within 3Å as condensed ions/atoms A around atom/molecule B. For condensed metal ions around the negatively charged Glu amino acid, the distances between the metal-ion and Glu residue averaged over last 10ns simulation data, 2.42 and 2.57 for NaCl and MgCl_2_ respectively confirm the effective larger size of K^+^ than Na^+^ ion. So not only the extent of ion-pair interaction also the smaller size of the Na atom than K may cause a pronounced interaction of Na^+^ ion with amino acid Glu of peptide over K^+^ ion as small cations are likely to bind on the surface of charged polypeptide. A comparison of the ratio (*t_c_*) of condensed cations around Glu and metal ions unbound to residue Glu as a function of time is shown in Figure 9. In Figure 9, the increase in the interaction of Na^+^ ions with Glu with time is an indicator of the influence of free Na^+^ ions within bulk-water on the growth of aggregates where by the interaction is constant on average in KCl case. The average number of condensed cations and anions for various cases are listed in Table II. Table II shows aweaker interaction of Cl ions with protein heavy-atoms.

**FIG. 9.**
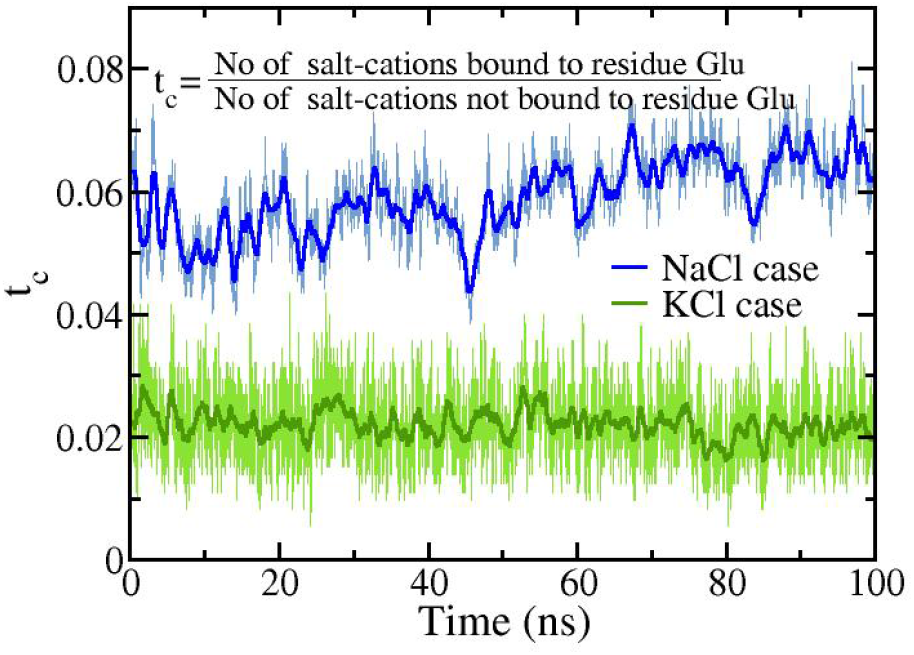
The ratio (*t_c_*) of the no. of salt-cations (Na^+^ and K^+^) bound to Glu and no. of salt-cations not bound to Glu in simulation as a function of time.

**TABLE II.** Number of condensed ions averaged over last 10ns MD data.

| Salt | Mean aggregate size ($\bar{S}$) | Cation-Glu<sup>a</sup> | CLA-peptide<sup>b</sup> |
| --- | --- | --- | --- |
| NaCl | 51.167 | 43.9105 | 0.78 |
| MgCl<sub>2</sub> | 27.62 | 4 | 1.812 |
| KCl | 19.18 | 15.68 | 0.638 |

<sup>a</sup> Average number of condensed salt-cations around Glu residue.
<sup>b</sup> Average number of condensed $\text{Cl}^-$ ions around heavy atoms of peptide.

Further the primary free-energy contributions of the peptide atoms that lead the polypeptide system toward forming the aggregates have been discussed in next section.

### D. Electrostatic, vdW interaction energy and solvation free energy of aggregates

Aggregation of peptides is usually considered to be a result of absence of repulsive electrostatic forces, the deceasing effect of solvation free energy of peptides and short-range of vdW interaction between peptides^99^. The decreased effects of solvation are attributed to the absence of H-bonds with water molecules and the formation of interpeptide H-bonds. Here, we estimate the binding energy between the peptides using Molecular Mechanics Generlised Born Surface Area (MM/GBSA)^100^ method. According to MM/GBSA the binding free energy (Δ*G_bind_*) of the peptide molecules to form aggregates can be estimated from the following sum,

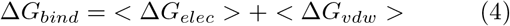

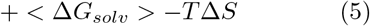

where the first two terms are standard molecular mechanics (MM) contributions originating from electrostatic and van der Waals interactions of aggregates, Δ*G_solv_* is the total solvation free energy obtained from the polar (Δ*G_polar,elec_*) and non-polar (Δ*G_np_*) contributions, and *T*Δ*S* corresponds to the entropic contribution. Here, non-boned interaction energy (vdW energy and electrostatic energy) of peptide molecules and electrostatic solvation free energy (Δ*G_polar,elec_*) have been computed by ‘namdenergy’ plugin of VMD. The nonpolar solvation energy (Δ*G_np_*) can be estimated from a linear relation to the solvent accessible surface area (SASA)^101,102^ given by,

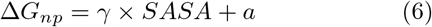

where γ and a are constants. In our analysis SASA has been calculated using 1.4Å as the effective radius of an atom with VMD, and the constants γ and *a* were set to 0.005 kcalmol^−1^Å^−2^ and 0.0 respectively^103^. The entropic term (TΔS) can be estimated by a normal-mode analysis of the vibrational frequencies^104,105^. However we have ignored the entropy contribution owing to the translational, rotational and vibrational degrees of freedom in our calculation due to its high computational demand. Finally the aggregation process proceeds via the counter-play between three types of contributions (Δ*G_elec_*, Δ*G_vdw_*, Δ*G_solv_*) of Δ*G_bind_*.

Figure 10(a) shows the evolution of electrostatic interaction energy calculated for the aggregates including the condensed ions around aggregates. The contributions originating from electrostatic interaction reveal that for NaCl salt solution systems, the net electrostatic energy is attractive in nature despite the presence of negatively charged Glu residue in each the peptides which should result in repulsive interaction between the peptides. The electrostatic attractive interaction between the salt-cations and Glu residue initiates the aggregation events by suppressing the effect of thermodynamically unfavourable electrostatic repulsive interaction between peptides. As noted before, for NaCl solution systems, a relatively higher number of Na^+^ ions are buried within the peptide aggregates compared to both MgCl_2_ and KCl solution systems. The snapshots of buried alkali-ions (monovalent) within the aggregates at the end of 100ns simulation are shown in Figure 10(b-c). The buried cations effectively renormalize the overall charge of the peptides, albeit in different ways (see next section), resulting in effective attractive interaction energies between peptides (in addition to short range hydrophobic energies).

**FIG. 10.**
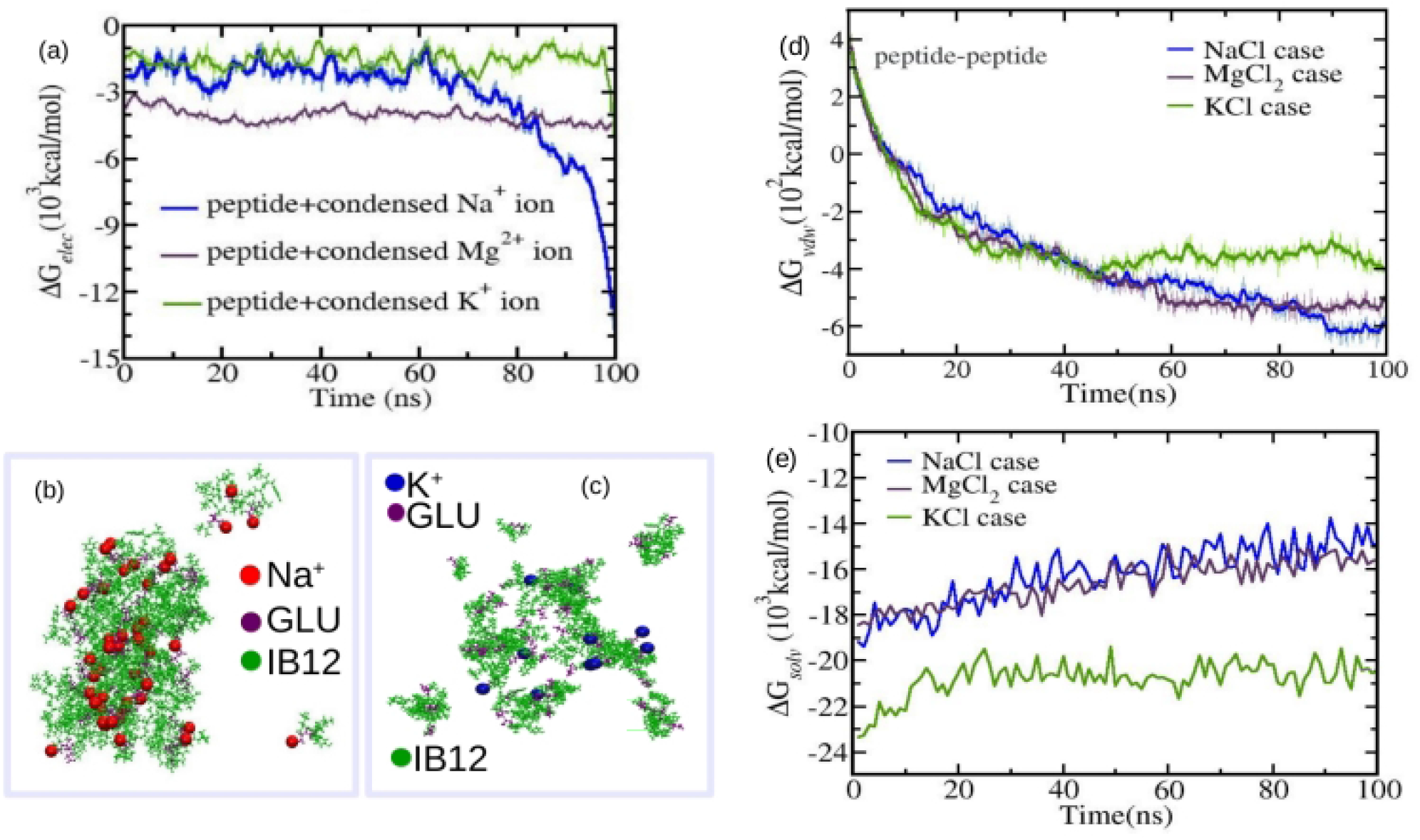
(a) Time-dependent comparison of electrostatic interaction energies for peptide aggregates and ions bound to peptide for three different IB12-salt systems. (b)-(c) Snapshots of buried alkali ions of salt within the IB12 peptide aggregates in salt solution captured at the end of 100ns simulation. The color code red, blue, purple and green present the Na^+^, K^+^, residue-Glu and peptide atoms (in CPK representation) respectively. Water molecules and Cl^−^ ions are omitted for clarity. (d) VDW interaction energy between the peptide atoms for three different IB12-salt systems. (e) Solvation free energy (G*_solv_*) of peptide molecules originating from the interaction btween water and peptide atoms.

In Figure 9 the number of Na^+^ ions bound to Glu was found to be increased with time indicating the cumulative nature of process, *i.e*. the propensity of Glu residue on the surface of aggregates to imbibe more Na^+^ ions from bulk water as the aggregate size increased and in effect this minimize the surface charge density which in turn further assists the growth of aggregates by weakening the effect of negative charge of peptides. Due to the less propensity of K^+^ ions to interact with Glu, K^+^ ions could not compensate the electrostatic repulsive energy of peptide molecules and cause small aggregates formation instead of a single large aggregate. Also, in Figure 10(c)-(d) the decrease in solvation energy term with time and increase in vdW energy with time respectively favor the aggregation process for all cases. We notice vdW energy is more negative for NaCl case than KCl case and that is because of larger aggregate size. As there is no significant interaction between the Mg^2+^ ions and charged peptide molecules, the Δ*G_solv_* contribution originating from hydrophobic interaction between the peptide molecules is the main driving force for preferential association of charged polypeptide in the presence of MgCl_2_ salt. Importantly, the electrostatic interaction of Na^+^-Glu and hydrophobic interaction both make favorable contributions to the formation of the large aggregates in NaCl-salt solution while unfavorable interpeptide electrostatic interaction and the solvation effect results the formation of small sized aggregates in KCl-salt solution.

### E. Charge renormalization of aggregates by different cations

It is clear from the above section that the final aggregate sizes are the result of the balance of three types of competing interactions namely protein-ion, ion-pair and water-ion interaction. This suggests that at certain values of aggregate sizes, the polypeptide system can achieve minimum energy and get stablized via the resultant interaction energy and results in discretization of aggregate sizes as obtained in the Figure 2. In this section we explore the effective renormalization of the peptide charge by the inclusion of salt, as compared to the control case, to understand the faster aggregation rates, in general, for salt-solutions compared to control system. Previous MD studies and electrophoretic measurements of the binding of divalent cations to insulin in aqueous salt solutions^106^ reported a significant reduction of the absolute value of the electrophoretic mobility of charged insulin in the presence of divalent alkali cations via charge rescaling. Previous works have also shown that multivalent ions can lead to overcompensation of charges, in addition to charge neutralization, and this can be a mode of aggregation of charged polymers such as DNA in the presence of multivalent ions^73–75^.

In Figure 11, a comparison of the effective charge of each peptide as a function of distance (*r*) from the peptide for the three IB12-salt solution and control system is shown. A striking feature of the plot is that compared to the control system, the effective charge is renormalized at much shorter distances in the presence of salt, though there are significant differences between the salts themselves. This strongly underscores the faster kinetics of aggregation formation of the charged peptides in the presence of excess salts. However, significant differences do exist among the charge renormalization behavior between the three salts at very short distances, which can be the reason behind different sizes and stabilities of the aggregates for the three salt solutions. For NaCl case, it can be seen that the charge neutralization is achieved at a very small distance from the peptide and this underpins the fastest and the most abundant aggregate size formation for NaCl salt solution. For both KCl and MgCl_2_ salt solutions, the analysis strongly suggests that the cations are not as effective as the Na cations in renormalizing the charges at short distances. Infact for Mg^2+^ ions, a charge inversion is clearly observed, where the net effective charge changes sign and this is consistent with previous studies on effects on multivalent cations on charged biopolymers such as DNA and other macroions^42,73^. In our analysis we also find that Mg^2+^ ions are prone to bound with Cl^−^ ions in water largely. Therefore, the polarization field of MgCl_2_ in water may be the cause of the observed charge inversion of charged peptide and that in turn may effect the aggregation rate. Indeed, for MgCl_2_ salt Mg^2+^ ions accumulate in the spherically symmetric surface only untill the point of neutrality is reached. Also, for MgCl_2_ salt, the electrostatic shielding effect of excessive number of Cl^−^ ions may cause a hinderance for Mg^2+^ ions to come close to peptide atoms.

**FIG. 11.**
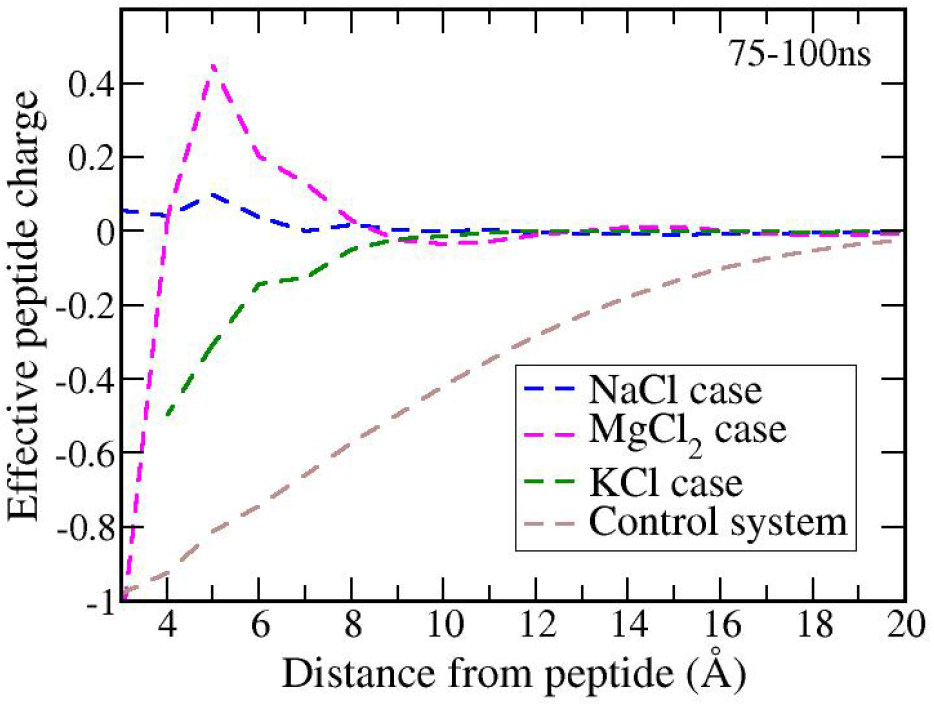
Effective charge of each peptide as a function of the distance from the peptide heavy atoms (i.e. sum of the net peptide charge together with the charge of all ions located within a given distance of the peptide averaged over all 64 peptides), as obtained from last 25ns simulation with NaCl (blue), MgCl_2_ (magenta), KCl (green) and control system (brown).

### F. Condensed ion lifetime distribution

Solvated ions have important functional roles in influencing the structure and dynamics of biomolecules^48–54^. In this context, lifetime measurement of ions i.e. the residence time of ions in the vicinity of peptide chains is an important tool in our understanding of the ion-protein interaction and subsequent insight into the effect of salt on the aggregation process of multipeptide system. Here we calculate lifetimes of condensed ions and report the distribution of the same as shown in Figure 12. An ion is considered to be a condensed ion, if it is at a distance <=3Å from any of the peptide atom. Figure 12 shows the distribution of lifetime on a log-log scale for cations and anions of salt separately. In Figure 12(a) we observe a change in the slope in the life time distribution plot of condensed cations, obtained from 100ns long IB12 trajectories, before and after 100ps time scale. The slope of frequency vs. time is nearly constant at the trailing part of the curve albeit small fluctuations. Since there may always be few ions which are likely to interact with peptides at any point of time, the frequency of lifetime distribution for such ions lies in region between 0-100ps of the timescale which is relaxation timescale of the solvated system. The interaction of ions with lifetime less than 100ps are removed from our consideration due to their strong interaction. The plot with greater than 100ps timescale indicates power-law behavior. When slope of the plot is high, the number of condensed ions with large lifetime is smaller than the number of condensed ions with low lifetime. In Figure 12(b) We don’t observe any significant difference in the slope of the lifetime distribution curve of condensed Cl^−^ ions for three types of salts, only difference is the increased frequency value of condensed Cl^−^ ions for divalent salt compared to monovalent because of higher number of Cl^−^ ions in simulation box in the case of MgCl_2_. Therefore, we can say that salt-anions don’t have important role in aggregation process of negatively charged IB12-salt system due to electrostatic repulsion between Cl^−^ ion and amino acid Glu residue. In the case of condensed cations the slopes of the histogram plot are different for three different types of salt. The slope of histogram plot of lifetime is less steeper for NaCl case than KCl and the residence time of +ve condensed ion around peptide atoms extends upto 10ns and 1.5ns for Na^+^ and K^+^ ions respectively, suggesting the higher residence times, hence higher interaction strength of Na^+^ ions with peptides than for K^+^ ions. In case of MgCl_2_, the number of Mg^2+^ ions near the peptides is small, and the initial Mg^2+^ ions get trapped within the aggregate at the very early stage of simulation and remain attached to the surface of amino acid Glu throughout the simulation and this results in a single data point near the 100ns time scale on the plot. On average, the Na^+^ ions show significantly greater probability to be bound with negatively charged polypeptide molecules in peptide-salt solution compared to other two cations.

**FIG. 12.**
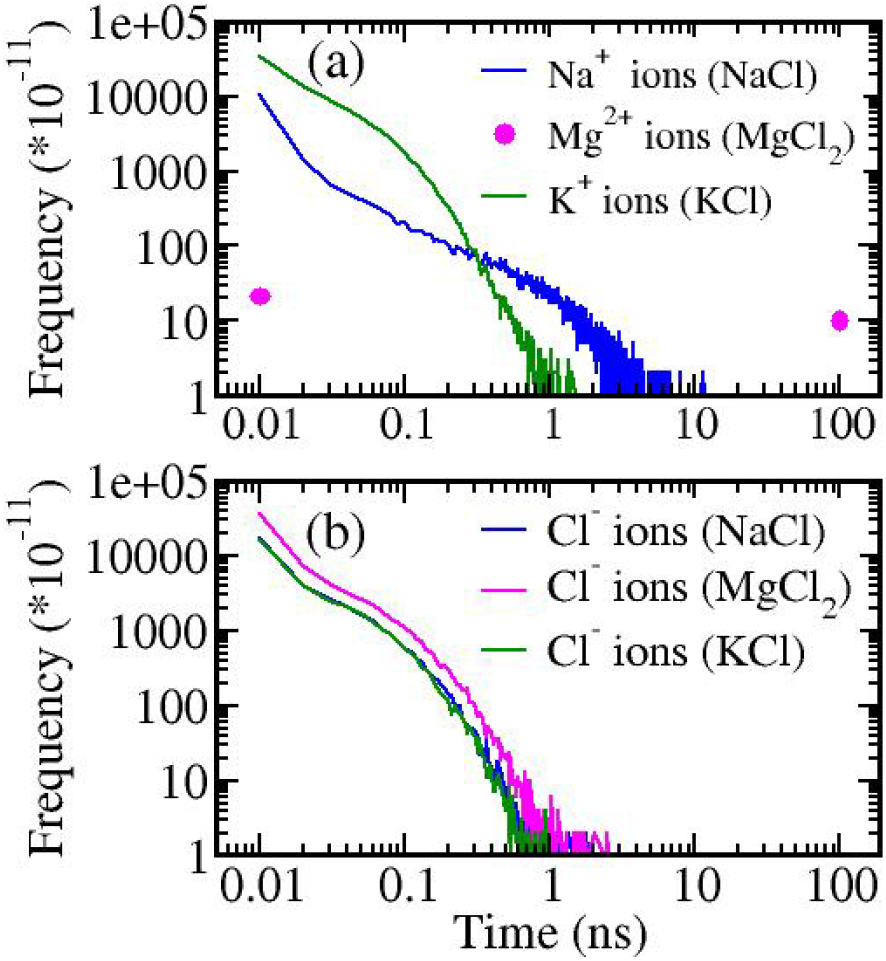
Historgram of condensed ion lifetime (*i.e*. residence time of an ion within 3Å of the backbone atoms of proteins) plotted on a double-log scale for cations(a) and anions(b) separately in three different salt solutions.

### G. MSM analysis of Aggregate size distribution

Figure 13(a)-(b) shows the evolution of fractional populations of *N*-mer aaggregates with propagation step (*n*) for top ten high-populated aggregate states in the case of NaCl and MgCl_2_ respectively computed using MSM analysis. The components of the transition probability matrix (*P*) were calculated as the fraction of *i*-mers undergoing a transition to a *j*-mer at a given propagation step. We observe that the system starts to equilibrate after 75ns, hence in our MSM analysis we used the total 100ns of two MD trajectories for each case for constructing the transition probability matrix.

MSM analysis for NaCl salt revealed that the top ten highly populated amorphous aggregate states are 52, 53, 54, 51, 55, 50, 58, 56, 57, 40-mer and the probability of these aggregate sizes increased with time and saturates after a certain propagation step. The aggregate of size 52 has the highest possibility to be formed for a 64-mer IB12 system with NaCl salt and the probability of higher order aggregate is decreased with size of aggregate. For the medium size cluster of size 40 we observe a slight bump on the occupancy of *N*-mer aggregates with a decrease in fraction indicating that the medium clusters were growing to larger amorphous aggregates (> 50-mer). However, in the case of MgCl_2_ salt, the smaller and medium size aggregates have higher fraction in the trajectory. Initially the small size aggregates (4-mer, 5-mer, 6-mer, 18-mer) have large fraction and they rapidly accumulate to medium size clusters (32-mer, 33-mer, 42-mer, 43-mer). The fraction of medium aggregates consolidates with propagation step but the formation of large aggregates was found to be less probable for MgCl_2_ salt solution compared to NaCl salt solution. This strongly suggests that at the initial stage of aggregation pathway, the smaller size clusters play key role in the formation of larger clusters and different salts can effect this differently.

**FIG. 13.**
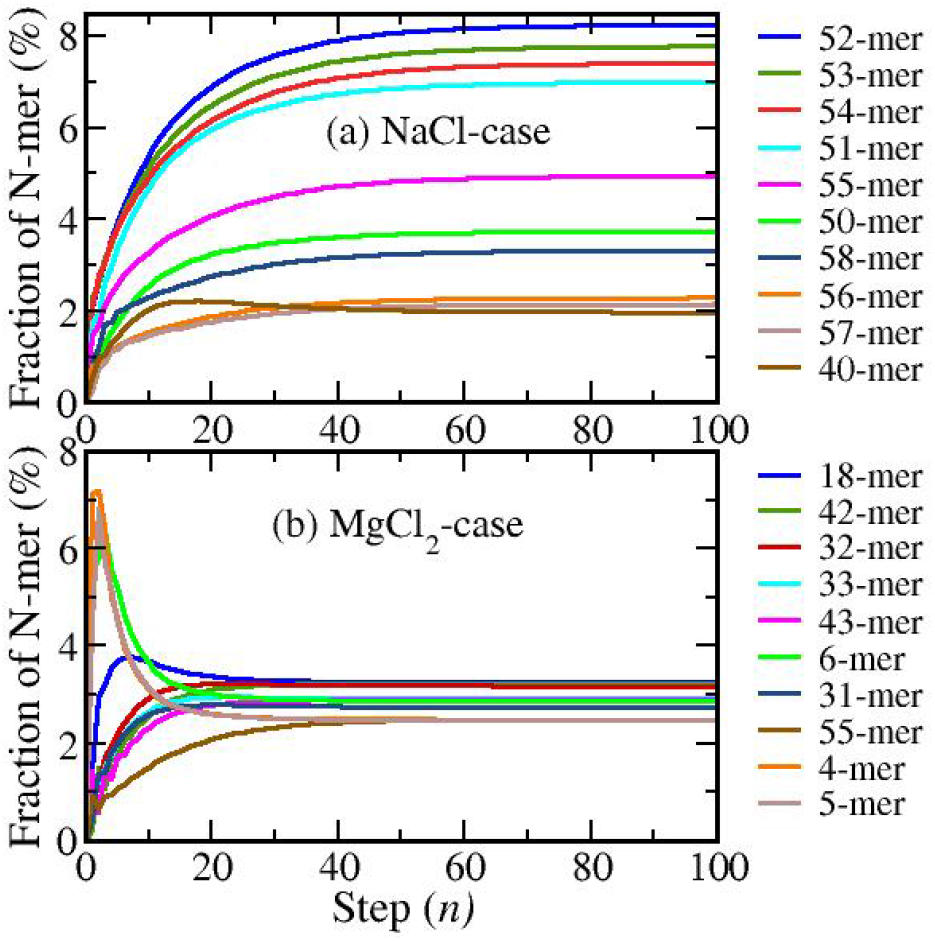
Markov State Model analysis of N-mer fraction for top ten higher populated aggregate states for NaCl(a) and MgCl_2_(b) salt solutions, computed by solving master equation.

## IV. DISCUSSION AND CONCLUSION

In this article, a systematic analysis of the effect of salt at high concentration on the amorphous aggregation of charged peptides (IB12) in water have been reported using MD simulations and detailed analyses involving MM/GBSA and charge renormaliztion by salt-ions. Two independent 100ns MD simulations with two monovalent salts (NaCl and KCl) and one divalent salt (MgCl_2_) corresponding to IB12-salt solutions, and a control simulation with no additional salt (except to achieve charge neutrality) are performed. Efforts have been made to obtain an understanding of the aggregate morphology of the multi-peptide system and the effects of not only the valency of the added salt cations, but also type of cation as well.

The results of this study strongly suggest that the presence NaCl salt in solution leads to the formation of largest aggregate of 64-IB12 peptides, compared to other salt systems studied, including another monovalent salt KCl. For MgCl_2_ salt, the simulations suggest that though the propensity to form large aggregates is higher than KCl, the stability of formed aggregates is weaker than that of NaCl salt systems. Possible reasons for this include the cation-peptide interactions, cation size, competing favourable electrostatic and hydrophobic interactions and effective charge renormalization capabilities of the cations considered. The simulations suggest that the formation of large aggregate in NaCl-salt solution originated from both attractive electrostatic interaction of Na^+^-Glu (negatively charged) and subsequent hydrophobic interactions between amino acids of peptide. However, in the case of MgCl_2_-salt solution, the predominant favourable interactions in aggregate formation are hydrophobic in nature.

Charge renormalization by the counterions has been suggested to be one of the important pathways of aggregation of highly charged polymer systems^73–75^. In these systems, the condensed counterions effectively render the similarly charged polymers neutral, thereby inducing aggregation via short-range attractive interactions^74^. The effective charge of each peptide (averaged over 64 peptides) as a function of distance from the peptide is measured and it shows strong indications that the presence of NaCl effectively neutralizes the system, which is not the case for the other two salt solutions considered in the study. Significantly, the absolute value of effective peptide charge is positive for MgCl_2_ case and indeed, the overscreening of peptide charge by the Mg^2+^ cloud around the aggregates drives the peptide aggregation in MgCl_2_ solutions. So it is worth to conclude that though final aggregate sizes for NaCl and MgCl_2_ are comparable, the mechanism of aggregate formation is fundamentally different for the two cases. In case of other monovalent salt, KCl, the smaller and unstable aggregate formation is due to the lack of interaction between K^+^ and Glu residue of the IB12 peptides. This leads to unfavorable inter-peptide electrostatic repulsive interactions, which possibly cannot be compensated by the favourable inter hydrophobic interactions and thereby results in smaller and more unstable aggregates. The calculation of lifetime of condensed ions also revealed that Na^+^ ions have much more propensity to be bound to the aggregates for much longer time than K^+^ ions. Markov state model analysis suggested that small aggregates consisting of 4~6 peptides accumulate at initial stages of aggregation and then they merge into large aggregates. Interestingly, our simulations for IB12 peptide in explicit solvent environment using recent all-atom CHARMM36 force field have displayed precipitation of peptide monomers into large amorphous aggregate state but do not show significant conformational conversion of soluble peptides into insoluble amyloid-like fibrils contrary to expectations from previous simulations^59,60^. It has also been recently suggested that the force field may play a role in elucidating the final conformations of such peptides in solution^107^. Therefore, caution is needed to chose appropriate force-field parameters in the protein aggregation studies. However, the primary focus of this study is not on the fibrillar formation of either single or multiple aggregates, but to understand the role of salts on the aggregation dynamics of charged peptides in solutions. Much longer simulations and multiple simulations are most likely required for detailed understanding of the effect of different force fields and their inherent propensity for certain secondary structure formations, which is beyond the scope of the present study.

## SUPPLEMENTARY MATERIAL

See supplementary material for the normalized aggregate size distribution of charged poly-peptide system in NaCl and MgCl_2_ salt solution with TIP4P-Ew water model, the g(r) function for ion-pair and water-ion interactions, and snapshots of condensed anions and water molecules around a single cation.

## Acknowledgments

The simulations were carried out on the supercomputing machines Annapurna and Nan-dadevi at the Institute of Mathematical Sciences.

